# Freely foraging macaques value information in ambiguous terrains

**DOI:** 10.1101/2024.10.11.617791

**Authors:** Neda Shahidi, Zurna Ahmed, Yuliya Badayeva, Irene Lacal, Alexander Gail

## Abstract

Among non-human primates, macaques are recognized for thriving in a wide range of novel environments. Previous studies show macaque’s affinity for new information. However, little is known about how information-seeking manifests in their spatial navigation pattern in ambiguous foraging terrains, where the location and distribution of the food are unknown. We investigated the spatial pattern of foraging in free-moving macaques in an ambiguous terrain, lacking sensory cues about the reward distribution. Rewards were hidden in a uniform grid of woodchip piles spread over a 15 sqm open terrain and spatially distributed according to different patchy distributions. We observed Lévy-like random walks in macaques’ spatial search pattern, balancing relocation effort with exploration. Encountering rewards altered the foraging path to favor the vicinity of discovered rewards temporarily, without preventing longer-distance travels. These results point toward continuous exploration, suggesting that explicit information-seeking is a part of macaques’ foraging strategy. We further quantified the role of information seeking using a kernel-based model, combining a map of ambiguity, promoting information seeking, with a map of discovered rewards and a map of proximity. Fitting this model to the foraging paths of our macaques revealed individual differences in their relative preference for information, reward, or proximity. The model predicted that a balanced contribution of all three factors performs and adapts to an ambiguous terrain with semi-scattered rewards, a prediction we confirmed using further experimental evidence. We postulate an explicit role for seeking information as a valuable entity to reduce ambiguity in macaques’ foraging strategies, suggesting an ecologically valid way of foraging ambiguous terrains.

**Author Summary:** In a novel and ambiguous terrain lacking sensory information about the location or distribution of food, foragers obtain information by sampling. This process is crucial for animals thriving in a new habitat. We allowed free-roaming macaques to forage at their own pace in a controlled terrain to which they had limited prior exposure. Based on their foraging paths, we developed a computational model representing an individual’s drive for reward-seeking, information-seeking, or energy preservation. These drives were represented as superimposing maps from the forager’s perspective. We found that information-seeking, continuous exploration of unknown areas, is crucial for foraging under ambiguity. This finding is consistent with a theory suggesting that animals, specifically humans and other primates, seek information to reduce their uncertainty about the environment. Our study suggests that the statistical properties of primates’ random foraging patterns reveal their complex decision-making process, including adaptation to novel environments.

## Introduction

Foraging monkeys adopt a variety of strategies in environments with sparse food sources [1]. Some species adapt to familiar habitats by memorizing and revisiting high-yield food sources [2] or following seasonal diets [3]. Others, however, use more flexible strategies that allow them to forage across a wide range of habitats and adapt to unfamiliar terrains. Among non-human primates, macaques are particularly notable for their ability to thrive in diverse geographical environments [4]. Individual macaques can travel several kilometers daily [5], covering relatively long distances for primates. Some individuals, typically adolescent males, disperse from their birth group to join others or form a new group [6]. Their diverse habitats, large home ranges, and the potential for dispersal suggest that macaques frequently encounter an unfamilair distribution of food sources, a situation best tackled by exploration and seeking information. This raises the question of whether and how information-seeking influences macaques’ choices of where to forage in an unfamiliar environment.

### Decision-making under ambiguity or uncertainty

In situations of ambiguity, where there is a lack of information about the existing situation, or uncertainty, defined by the unpredictability of outcomes, macaques tend to explore unknown options or actively seek information. Comparative studies on gambling for food in the presence of ambiguous options have shown that macaques, along with gorillas, chimpanzees, and orangutans, recognize the potential rewards associated with these ambiguous choices [7]. Notably, macaques demonstrate a preference for mildly uncertain reward options over certain or entirely random ones [8], particularly when rewards are sufficiently available [9]. This information-seeking behavior is suggested to be driven by an intrinsic motivation to reduce uncertainty [10]. The evidence for this comes from studies showing that macaques choose informative options, whether or not the information directly enhances rewards in future choices [10], [11]. These findings raise the question of whether macaques’ foraging strategy under conditions of ambiguity or uncertainty encompasses targeted information seeking.

### Sampling uncertain or ambiguous options

Sampling uncertain food sources while foraging provides potentially useful information about the hidden structure of the environment. Exploration, formulated in reinforcement learning theory as randomness in the decision-making policy [12], is observed in many species when foraging stochastically refilling sources. The matching law [13]–[19] predicts that animals allocate their effort to each source proportional to its value, determined by its refill rate. However, many species tend to over-sample the low-rate source when the refill time is unpredictable, compared to the prediction of the matching law [13]. In macaques, this behavior was explained by a model of information gathering under uncertainty, enabling foragers to detect unnoticed fluctuations in the refill rate of the low-rate source [20].

Searching space for ambiguous food sources has been conceptualized in patch-wise foraging, where the forager decides between staying in the current habitat or relocating to a new one. Theoretical works predict that even an optimal forager, who maximizes the future rewards, under-samples a high-yielding patch and vice versa [21]. Patch leaving decisions were also explained as an evidence accumulation process in which each sample is a noisy source of information about the structure of the foraging terrain [22]. This view on foraging is a physiologically plausible way to explain the inherent randomness in animal behavior when navigating an ambigious terrain, as we investigated further in our study.

While these studies explain how animals choose among a finite set of food sources, the manifestation of exploration and information seeking on a continuum of choices, for example, when foraging ambiguous terrains with scattered sources of hidden food, is largely unknown.

### Spatial search algorithms and heuristics

Foraging an ambiguous terrain may be conceptualized as a spatial search challenge, where the searcher sequentially samples potential locations for hidden food. Each sample may yield food or not, but in either case, it provides information on the food’s spatial distribution. An optimal search strategy may minimize the number of samples tested by excluding less probable locations, shortening the navigation path between sequential samples, or both. For instance, when sampling is effortful because it includes digging for buried resources, the number of samples may be minimized by choosing the location of the next sample adaptively after gaining information from the current sample [23], [24]. Conversely, when relocation is effortful, for example, in avalanche rescue using beacons with a known range, minimizing the traveled distance by systematically scanning the environment according to a predefined pattern becomes crucial [25].

While optimal search algorithms that carefully minimize the searching time and effort are conceivable for known properties of the environment (the range of the beacon in the example above), biological systems often rely on heuristics. For example, when searching for lost keys in an apartment, typically, people go through possible places in a semi-random pattern instead of scanning the apartment from one corner to the opposite corner. A seemingly random heuristic may not be optimal for a static and familiar foraging terrain. However, considering a terrain’s ambiguity, i.e., lacking cues about food distribution, as a prominent property of natural habitats, such heuristics are beneficial [26].

### Random walks for spatial foraging under ambiguity

When the distribution of potential food sources is uniform rather than patch-wise, for example, when a forager searches for hidden insects or seeds in a meadow, a forager is free to take steps of any size in any direction. Therefore, the decision process differs from a patch-wise search in which the forager makes binary choices between staying and leaving. A ubiquitously reported foraging path in land and sea animals is a scale-free pattern known as Lévy walk or flight [27], [28] in which the forager takes steps of any size in any direction, but the step size is chosen from a heavy-tailed probability density function. Simply put, a Lévy often travels short, sometimes medium-sized, and rarely large distances between consecutive searches.

Lévy walks might as well emerge from the distribution of potential food sources rather than forager decision-making. For example, the Lévy distribution of traveled distances between consecutive searches for spider monkeys in a forest was explained by the scale-invariant distribution of distances between trees [29], which in turn is affected by spider monkeys’ navigation pattern because they are one of the main seed distributors for their food sources [30]. Therefore, whether a Lévy-like pattern originates in the forager’s decision-making process or is dictated by the natural food distribution remains unclear. The use of uniform terrains, where the distribution of potential food sources does not bias foragers’ strategy, is crucial for understanding foragers’ decision-making process.

### Area-restricted search

Although a Lévy-like foraging pattern effectively balances energy preservation with information-seeking, it does not explain alterations in forager’s paths when encountering food-rich locations [31]. For example, gophers excavate more tunnels in areas with high densities of their favorite plants than in areas with low densities [32]. Species of dolphins linger in sites of the ocean in which they have encountered prey minutes earlier [33]. This foraging strategy, known as area-restricted search, is particularly effective when the food distribution is localized or patchy (Motro & Shmida, 1995). Therefore, while a Lévy-like random walk balances energy preservation and exploration, an area-restricted search adjusts the foraging path to exploit a discovered food patch [31].

In some animals, finding food transforms the search pattern from *roaming* to *dwelling* [31]. For example, when *C. elegans* searches for food in a Petri dish, it swims in a relatively straight path (roaming). Once it encounters food, it substantially restricts its search to the nearby area by reducing its speed and moving in a convoluted path (dwelling) [34]–[36].

A substantial switch from roaming to dwelling indicates the end of exploration part of foraging and the start of exploitation [34]–[36]. However, humans and many other foragers balance exploration and exploitation throughout their search, meaning they never stop exploring [37]. While exploration entails random searches in this context, we aimed to understand a forager’s balance between targeted information-seeking and reward-seeking in an ambiguous terrain.

### Our approach

Here, we investigated macaques’ spatial foraging pattern on a two-dimensional ambiguous terrain, without sensory cues revealing the structure of food distribution. In our Exploration Room platform [38], we developed a controlled environment where male macaques freely foraged for hidden food within a spatially distributed, quasi-continuous pile grid. We video-recorded each session, identified the sequence of searched piles and their outcome, and identified the spatial structural characteristics of individuals’ foraging patterns. Our goal was to investigate how seeking information from an ambiguous terrain manifests in macaque’s foraging paths.

## Results

On the floor of our Exploration Room platform [38], four male macaques individually foraged for rewards in a grid of woodchip piles (Fig. 1A, B). This platform provided an experimentally controllable, distraction- and obstacle-free environment for foraging macaques. The grid consisted of 81 to 108 woodchip piles, each with an approximate diameter of 10 cm, arranged with a pitch of 25 cm (Fig. 1B). In each session, the reward pieces were invisibly hidden under 21 of the piles (*filled* piles), according to a pre-determined *abundance map* with a localized set of filled piles (Fig. 1B). The center of this set was selected randomly in each session in a way that intended locations of filled piles do not pass the edges of the pile grid.

**Figure 1.**
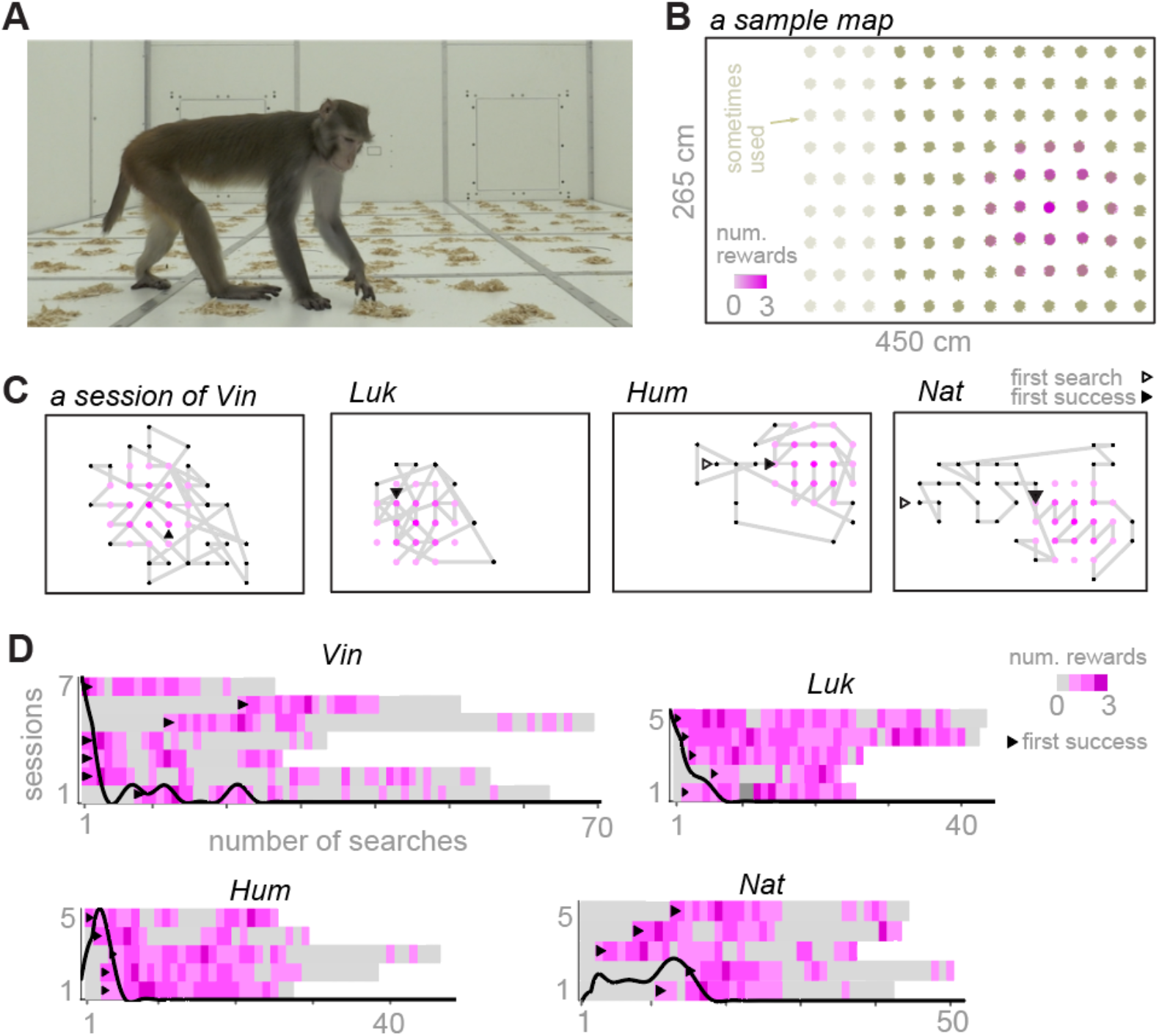
The experimental setup and statistical properties of the search patterns. **A)** Monkey Vin in a floor foraging session in the open arena of the Exploration Room. **B)** The grid of piles in the arena and an example of the hidden localized abundance map. The color intensity indicates the number of reward items hidden in a single pile, which was always 3 in the center and 1 at the outer margin of the circular disk, and zero elsewhere. **C)** Example foraging paths from each monkey (session #7 of Vin, #5 of Luk, #3 of Hum, and #5 of Nat). We let the animals stay in the room until they lost interest in foraging and spent more than one minute waiting in front of the exit gate or roaming without foraging. **D)** Raster of foraging outcomes in each session. The first successful pile search is marked with a triangle. The probability distributions of the first successful pile search for each monkey overlap the rasters.

All monkeys searched the terrain stochastically, starting from a seemingly random pile and navigating most of the terrain. Across all sessions (7 sessions of monkey Vin, 5 sessions of Luk, 5 sessions of Hum, and 5 sessions of Nat), monkeys found at least 42% and at most 100% of the filled piles. We kept the number of sessions small to probe the animals’ foraging strategy independent of long-lasting experience, emulating foraging in a novel ambiguous habitat. The first filled pile was found at the earliest in the first search and the latest after 22 searches. Accidentally finding a filled pile at the first attempt was unsurprising, given that the filled piles accounted for 19% (21/108) to 26% (21/81) of the grid.

### Patch-wise or Lévy-like exploration

We investigated seemingly random foraging paths to understand whether they resemble any of the observed spatial foraging patterns in natural habitats. Food sources, such as vegetation in natural terrains, may be distributed patch-wise or fractal-like (Fig. 2A *left and middle*), potentially biasing a forager’s path toward patch-wise or Lévy-like patterns. Instead, a uniform distribution (Fig. 2A *right*) allows a range of random search patterns. Hypothetically, the distribution of step sizes, defined as the Euclidean distance between locations of consecutive searches, reveals differences between patch-wise, characterized by a bimodal distribution representing within patch steps in one peak and between patch steps in another, and Lévy-like, characterized by a long-tailed distribution (Fig. 2B). A Brownian random search, not reported as animals’ foraging path but a viable alternative random walk, is characterized by a narrow range of step sizes, represented as a cropped Gaussian distribution [39] (Fig. 2B).

**Figure 2.**
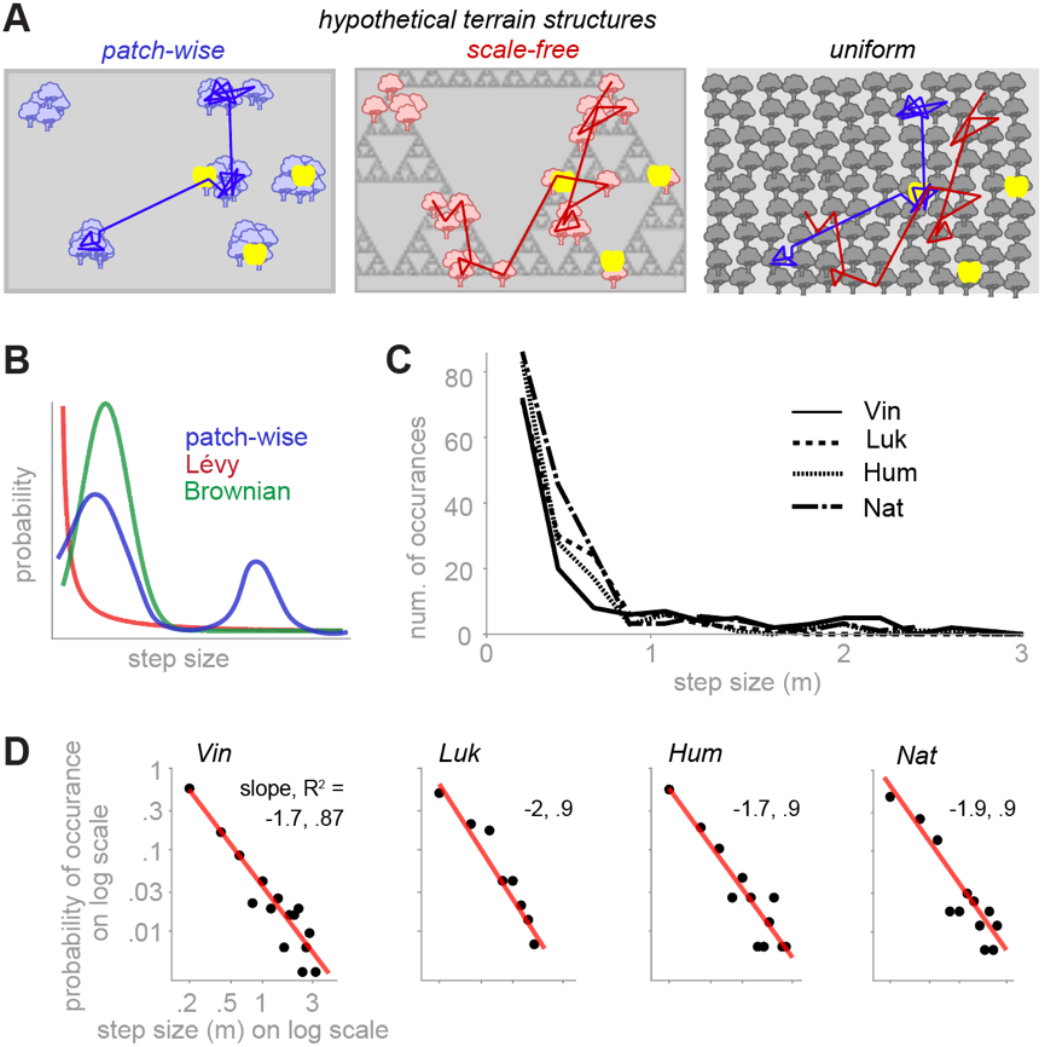
Step size distribution was closer to a Lévy-like random walk than a Brownian or patch-wise distribution. **A)** Three hypothetical types of terrains with a patch-wise (left), a fractal-like (middle), and a uniform (right) distribution of potential food sources. Three yellow dots show locations of hidden food pieces, which, in theory, could be identical across many distributions of potential food sources, among which three are shown here. This illustration suggests that when food is hidden, the spatial pattern of foraging may be biased toward patch-wise or Levy-like exploration merely by the distributions of potential food sources, independent of the actual location of the hidden food. In contrast, the uniform distribution minimizes this bias because it equally allows patch-wise (blue) and Lévy-like foraging (red). **B)** Hypothetical distributions of step sizes for a patch-wise, a Lévy-like, or a Brownian walk. **C)** The frequency of occurrences of step size, pooled across sessions of each monkey, binned into 20 cm bins. All distributions were unimodal (p = 0.09 (Vin), 0.067 (Luk), 0.13 (Hum), and 0.07 (Nat); Hartigan’s dip test). **D)** same as B but shown on double log scales and normalized as a probability density function.

We quantified step sizes by calculating the Euclidean distance between consecutively searched piles. Although we occasionally observed patterns resembling a patch-wise search (Fig. S1), the distributions of the step sizes, pooled across all sessions of each animal, did not indicate bimodality (Hartigan’s dip test, p > 0.05) and were heavy-tailed (Fig. 2C), suggesting a lack of evidence for a patch-wise search. We calculated the probability of occurrence, i.e., the normalized frequency of occurrences, for 0.2 m step-size bins. We tested whether it decreases as a power law function [27] of the step sizes, which generates a linear decrease on a double-logarithmic scale (see Methods). In contrast, a Brownian walk is expected to create an inverse bell shape on a double logarithmic scale. For all monkeys, the probability of occurrence fits a linear function (Fig. 2D) with a slope that falls within the range [-1, -3], which is considered a signature of Lévy-like foraging [27]. Therefore, although monkeys frequently chose nearby piles, they explored the grid by flexibly choosing new locations at large distances within the grid’s boundaries. Observing Lévy-like, rather than bi-modal step size distributions in our dense and uniform terrain, designed to minimally bias foraging choices, suggests that binary choices between staying nearby piles or exploring far piles were not behind macaques’ foraging strategy.

### Less exploration after finding food?

We investigated whether food encounters alter the animals’ between-pile step sizes or foraging paths. We compared each monkey’s step size after a success, i.e., after encountering a full pile, to a failure, encountering an empty pile. All monkeys, on average, chose a nearer pile (smaller step) after a success compared to a failure (Fig. 3A), suggesting that they expected to find more rewards in the vicinity of discovered rewards. Although such expectation sounds reasonable for a macaque whose experience with food collection is from natural sources such as trees, bushes, and ant insect nests, it is not immediately obvious for a lab macaque. Because monkeys could learn over time that the filled piles were clustered together, the difference between the averages of post-success and post-failure step sizes could have come from the late sessions of each monkey. However, the separability of the post-success and post-failure step size distributions did not reveal a trend across sessions of each monkey (Fig. S2), suggesting that staying in the vicinity of discovered food was not learned from the structure of the hidden food abundance map.

**Figure 3.**
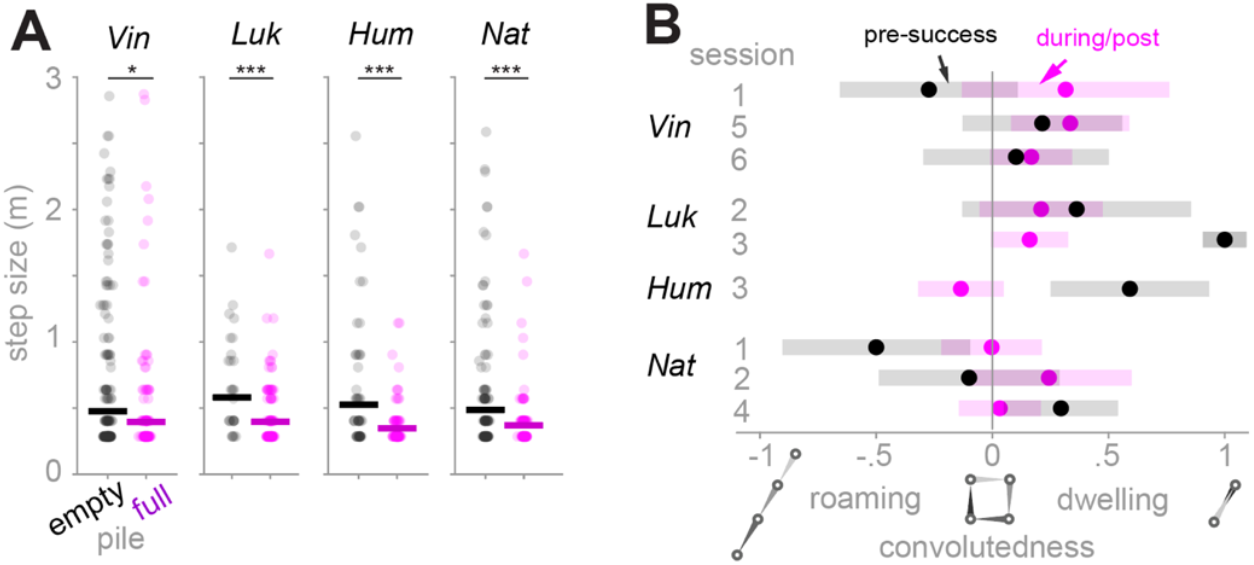
Monkeys favored, but not restricted to, searching piles in the vicinity of discovered rewards. **A)** Step size in meters after encountering an empty or a filled pile. Pile searches were pooled across sessions of each monkey (success/failure = 135/189 (Vin), 114/33 (Luk), 90/58 (Hum), 94/108 (Nat)). The geometric mean of step sizes after encountering empty versus filled piles is shown with a horizontal line. For each monkey, the geometric mean of step size after encountering filled (purple) piles was smaller than empty (black) piles (p = 0.02 (Vin), 0.006 (Luk), <<0.0001 (Hum), and 0.001 (Nat); Wilcoxon rank-sum test with FDR multiple comparison correction). **B)** For each session with at least 4 pile searches before and after finding the first reward, a convolutedness index (see methods) was calculated for the pre-reward sub-path, defined as the sequence of pile searches before encountering the first reward, as well as the during/post reward sub-path, defined as the sequence of pile searches including and after encountering the first reward. The convolutedness averaged across all sub-paths was slightly positive (0.2; p = 0.03, Wilcoxon signed-rank test). However, the pair-wise difference between the pre- and during/post-reward sub-paths was not significant (p = 1, Wilcoxon signed-rank test). Shaded bars represent the standard deviation of the convolutedness for sub-paths of length 4 (gray) or 8 (magenta).

Next, we asked whether encountering food switched monkeys’ foraging path from roaming to dwelling. We define roaming in our context as moving on a more or less straight line, while dwelling is characterized by many turns in short distances [31]. We defined a *convolutedness* index as the sum of the projection of each step on the previous step, normalized by the length of the sub-path (see Methods). This index is -1 for a straight path, 0 for a path with only right angles, and 1 for a path in which each step is the reverse of the previous step (lapses). By this definition, convolutedness is in [-1,0] for roaming and in [0, 1] for dwelling. For a sub-path passing via 4 piles, -1 indicates a straight alignment of visited piles, 0 indicates a square alignment, and 1 indicates repeated lapses between two piles (Fig. 3B).

For each session, we divided the foraging paths into pre-reward sub-path, ending at the first food encounter, and post-reward, starting from the first food encounter. For all sessions for which the length of the pre/post reward sub-paths > 4, the average convolutedness was positive (Fig. 3B). However, comparing the convolutedness of pre-reward sub-paths to post-reward sub-paths, we did not find a systematic shift from roaming to dwelling.

A lack of clear shift from roaming to dwelling due to finding a reward (Fig. 3B) might seem contradictory with the finding in Fig. 3A, where we showed that monkeys favored searching a nearby pile after encountering a filled pile. However, findings in Fig. 3A and B reveal a crucial difference between a roamer-dweller and a forager who continues to explore even after finding rewards. Essentially, these results point toward a foraging strategy in which the monkeys temporarily adjusted their step size to stay near the reward area but did not stop to explore. In theory, while roaming/dwelling patterns are optimal for a localized abundance map, continuous exploration allows the forager to adapt to a range of abundance map structures without needing to learn the map’s structure by prior experience.

### Seeking information in space: a kernel-based model

Searching the vicinity of discovered food, reported in the previous section, suggests that the forager assumes spatial continuity of the hidden food’s distribution. By revealing the content of a pile, full or empty, the forager gains information about the content of neighboring piles if the abundance maps have such spatial continuity and are not purely random. This makes an ambiguous pile a source of information extending to its neighborhood. Therefore, an area of the room with many ambiguous piles contains overlapping information sources, which is attractive to an information-seeker aiming to reduce the environment’s ambiguity. In principle, the spatial distribution of ambiguity adds a new dimension to a forager’s explorative decisions, besides reward-seeking and energy preservation, by encouraging the forager to gain information from ambiguous areas.

We sought to determine the effect of information seeking as an additional factor to reward-seeking and energy preserving, to explain animals’ foraging patterns. We simulated foragers considering these three factors to various degrees. Briefly, an agent chooses its next pile by sampling from a 2-dimensional probability distribution over the grid. We defined a map as a superposition of 2-dimensional Gaussian kernels centered at piles [40], [41]. In a virgin terrain, consisting of only unsearched piles, the *ambiguity map* (Fig. 4A, 1^st^ row, step 1) consists of the superposition of information kernels of all piles. Because the reward locations are unknown in this terrain, the *reward map* (Fig. 4A, 2^nd^ row, step 1) started empty. The *proximity map* (Fig. 4A, 3^rd^ row, step 1) consists of one proximity kernel around the forager’s current location on the grid. The probability map (Fig. 4A, 4^th^ row, step 1) is computed as a weighted sum of the information, reward, and proximity maps, normalized to have a sum of one. After each pile search, the ambiguity kernel of the searched pile is removed because the content of the pile is known (Fig. 4A, 1^st^ row, steps 2-20). If the pile contains rewards (for example, Fig. 4A, 2^nd^ row, step 10), the reward map is updated by adding reward kernels, one kernel per reward piece, centered at the searched pile. The proximity map is updated so that the proximity kernel is centered at the last searched pile, i.e., the current location of the forager (Fig. 4A, 3^rd^ row, steps 2-20).

**Figure 4.**
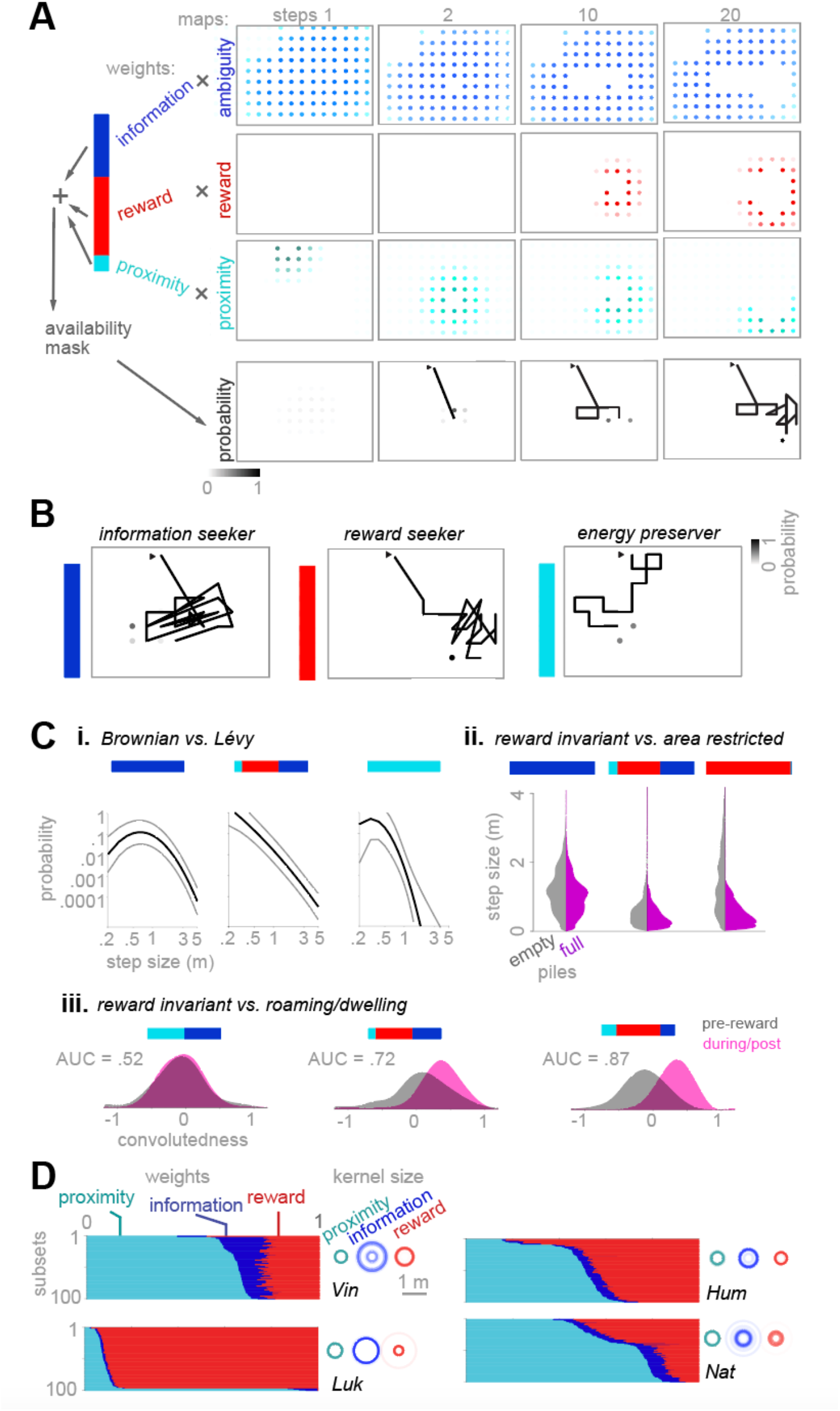
A kernel-based model of information, reward, and proximity explains the spatial properties of foraging paths. **A)** Simulation data from a generative model of spatial foraging in which at each step of foraging, the probability of choosing any of the available piles as the next pile is determined using a 2-dimensional probability map (see Methods). This map overlays many 2-dimensional kernels centered at ambiguous, rewarded, and proximal piles. The resulting path of a simulated agent on similar terrains to the experimental sessions was determined by the weighted sum of ambiguity, reward, and proximity maps. At each step of the simulation, the probability that each available is chosen as the next pile to search is determined as a weighted sum of ambiguity, reward, and proximity maps. Each map consists of a superposition of discretized kernels. The sizes of ambiguity, reward, and proximity kernels, defined as the standard deviation of the 2-dimensional Gaussian functions, were free parameters of the model. In the shown generative model, we used ambiguity kernels with standard deviations of 0.9 meters, reward kernels of 0.3 meters, and a proximity kernel of 0.6 meters. The ambiguity kernel’s standard deviation indicates that revealing a pile’s content provides information about up to 3 piles in each direction. The standard deviation of the reward kernel indicates that finding a reward is very likely in the adjacent piles of a filled pile. The standard deviation of the proximity kernel indicates the distance a monkey can reach by stretching its arm and body to reach a pile without relocating the full body. Regardless, similar simulation results were achieved using other values within reasonable ranges of these values. The next pile was randomly sampled from the 2-dimensional probability density function (4^th^ row). **B)** Simulation of three marginal agents: An information seeker (1^st^ row) chooses the pile at the peak of the information map. This agent moves to a new neighborhood of the map when the information in the current neighborhood is depleted, which occurs when one or more piles are searched. A reward seeker (2^nd^ row) moves randomly before encountering the first reward. After that, it stays within the vicinity of discovered rewards. An energy preserver (3^rd^ row) only takes the shorter possible steps in random directions, even after sampling empty piles. **C)** Selecting the composition of information, reward and proximity weights, the generative model prodices a variety of foraging strategies. i) a pure information seeker (left) or a pure energy preserver (right) produce Brownian-like random walks, as identified by an inverse bell share distribution of the step sizes while a balanced set of weights generates Lévy-like walks (middle). ii) A pure information seeker does not shortner step sizes immediately after encountering filled piles (left). The average of step size immediately after encountering a filled lowers as the weight composition transition into a balanced set (middle) or a pure reward seeker (right). iii) a reward invariant weight set (zero weight for the reward) produces the same distribution of convolutedness pre- and post-the 1^st^ reward encounter (left). Increasing the reward weight gradually shifts two distributions apart, indicating a roaming/dwelling strategy (middle and right). **D)** Fitting the model to the foraging paths of each monkey using maximum likelihood estimation (see Methods). 100 subsets of the pile searches, pooled across sessions (324, 147, 148, and 202 pile searches of Vin, Luk, Hum, and Nat), were used for training. Each subset contained 80% of the total number of pile searches. Left of each panel: The weight sets resulting from each training were sorted according to the value of the proximity weight for better visibility. Right of each panel: The fitted kernel sizes were shown as concentric transparent circles with the cicle’s radius showing 2 times the standard deviation of the Gaussian kernels.

Using this concept as a generative model, we generated an example forager weighing information, reward, and proximity in a balanced way (Fig. 4A), which roughly resembled the experimentally observed foraging paths. Using various sets of weights, foraging paths with diverse statistical properties emerge (Fig. 4B and C). For example, Lévy-like foraging emerged from a balanced weight set, while Brownian-like foraging (Gaussian distribution of step sizes) emerged from an information-dominant weight set (Fig. 4B, *left*; Fig 4Ci). Alternatively, a reward-dominant weight set strongly favored piles in the vicinity of discovered rewards, shortening the steps taken after reward encounters (Fig. 4B, *middle*; Fig. 4Cii). The convolutedness index for a reward- and proximity-dominant weight set switched from negative to positive after the first food encounter, indicating a switch from roaming to dwelling (Fig 4B, *middle*; Fig 4Ciii). Finally, a proximity-dominated weight set generated a crawling forager, strongly favoring piles adjacent to the current pile (Fig. 4B, *right*).

Comparing the results in Fig. 2 and 3 with the generative model in Fig. 4A-C suggests that monkeys balanced information, reward, and proximity weights in their foraging strategy. However, observing the foraging monkeys in our experiment, we expected individual differences not meaningfully explained by the results reported in Figures 1-3. For example, by qualitatively observing the behavior of the animals in the recorded videos, we expected Vin to be more explorative than other monkeys while Luk was more driven by reward or proximity.

We fitted the kernel-based model to each monkey by choosing the weights and the kernel sizes to maximize the likelihood of observing their foraging paths (see Methods). We pooled pile searches across sessions of each monkey and then fit the model to 100 randomly selected subsets of the pile searches from the pool of pile searches over sessions of one monkey (see Methods). The fitting results (Fig. 4D) revealed that while Vin weighed information higher than other monkeys, Luk weighed rewards more than others (Fig. 4D *left*; Fig. S3). Additionally, the size of the ambiguity kernel was typically larger than the reward and proximity kernel (Fig. 4D *right*). Therefore, ambiguous regions emerged from highly overlapping ambiguity kernels where most piles were unsearched. This result is consistent with the qualitative assumption that an information-seeking forager prefers an area of the terrain consisting of unsearched piles to a area in which some of the piles were searched and some not. The reward and proximity kernels were typically smaller, suggesting that proximity to rewards or self was defined as arm-reachable piles. All three types of kernels spanned a distance beyond the grid’s pitch, which was about 30 cm, suggesting that the monkeys have assumed spatial continuity in the hidden abundance map.

### Does the structure of the hidden abundance map matter

Fitting a model of spatial kernels to the foraging paths of monkeys provide evidence that they assumed spatial continuity in the hidden abundance map. This assumption happens to be true for our localized abundance maps with a large, single, localized set of piles containing hidden rewards. That raises the question of whether macaques’ assumption spatial continuity holds for abundance maps violating this assumption. To understand the effect of the continuity of the abundance map on foraging paths, we first simulated the forager from Fig. 4Cii *middle*, with a balanced proximity, reward and information weights, on terrains with scattered abundance maps (Fig. 5A). We found that for this type of abundance map, encountering a filled pile shortens the step size (Fig. 5B), equivalent to the localized abundance maps (Fig. 4Cii *middle*). Given that two of the monkeys, Vin and Nat, weighed rewards in a balanced way, as in the simulated forager in Fig 4Cii middle, we expected that they would shorten their step size on the scattered map as well.

**Figure 5.**
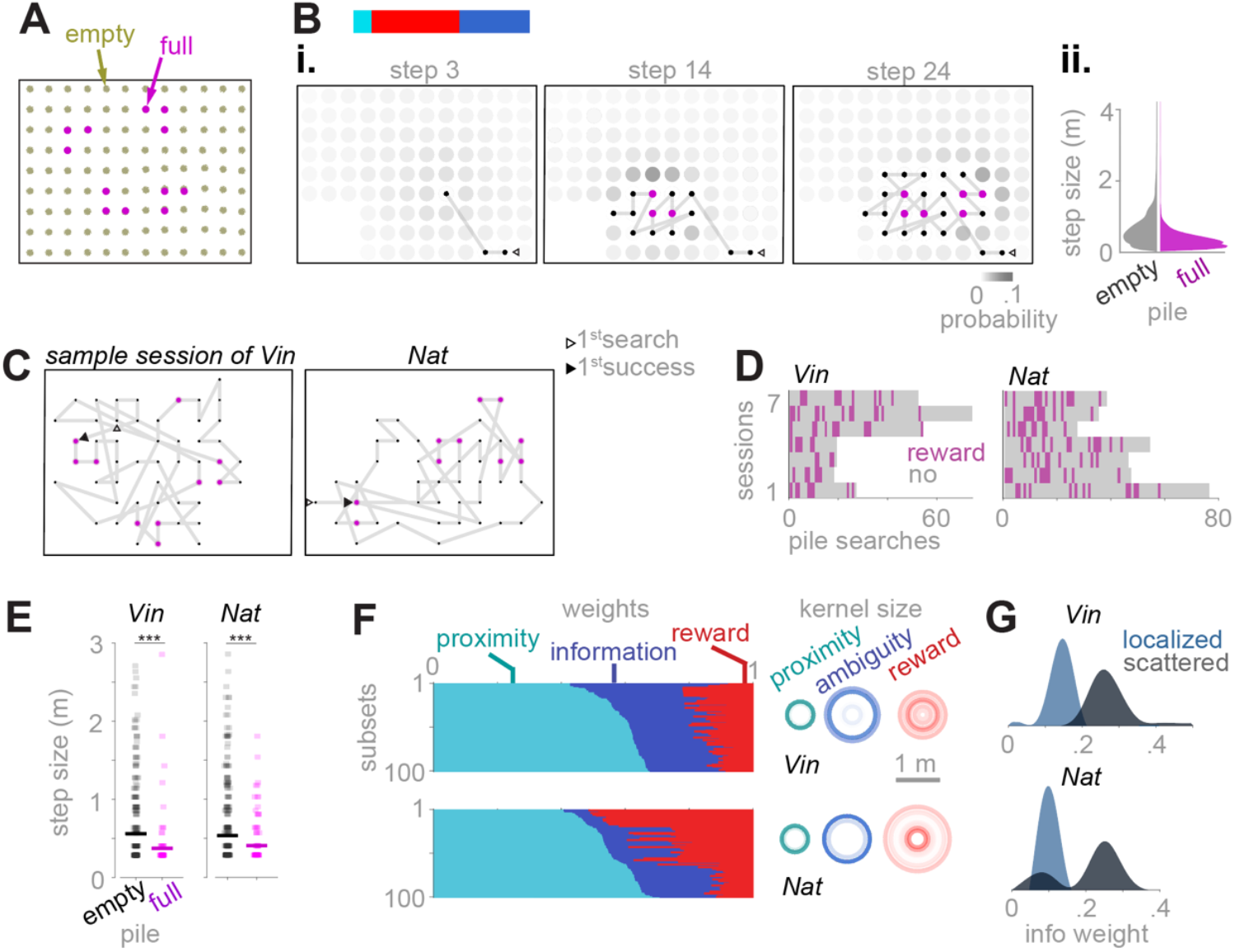
Simulated agents and monkeys favored the piles in the vicinity of discovered food on scattered maps. However, monkeys were more explorative compared to localized maps. A) A sample scattered map consisted of 4 groups of filled piles, each with three piles in a letter ‘L’ shape B) The simulated agent in Fig. 4Cii middle, here applied to scattered abundance maps. i) an example simulated foraging pattern. ii) the distribution of step size, after encountering an empty or filled pile, respectively. Results pooled across 100 simulated sessions. C) Example sessions of two monkeys on scattered maps. D) Reward raster of two monkeys, 7 sessions of randomly selected scattered maps for each monkey. E) Equivalent to Fig. 3A, but for the scattered maps shown in C. F) Fitting of the monkeys foraging patterns, equivalent to 4D, but for the scattered maps. G) Comparing information weights across localized and scattered abundance map structures for monkey Vin and Nat. For both monkeys, the information weights were higher for scattered maps (decodability, as the area under R.O.C., was 1 for Vin and 0.8 for Nat).

We tested Vin on foraging terrains with scattered maps, performing 7 consecutive sessions several months after performing the task with localized maps. Nat performed 7 sessions, which were randomly interleaved with 5 localized sessions (the results of Vin’s and Nat’s localized sessions were already discussed in Fig. 1-4). Despite this difference in the session arrangement, we found consistent pattern of encountering empty and folled piles for two monkeys (Fig. 5D).

Similar to Fig. 3A, encountering a filled pile shortened the step size (Fig. 5E), suggesting that monkeys assume spatial continuity of reward distribution. The average step size across scattered sessions was comparable to the localized sessions (Vin: 0.64 m for localized maps and 0.72 m for scattered maps, Nat: 0.61 m for localized maps and 0.69 for scattered maps; unpaired *t*-test, p = 0.21 for Vin and 0.20 for Nat).

Restricting search to the vicinity of discovered rewards is particularly useful when the abundance map is localized or patchy [42]. This strategy is expected to yield success on our localized maps more frequently than on our scattered maps. Unexpected failures, for example, when not finding rewards in the vicinity of discovered rewards, are proposed to generate internal error signals, leading to an adjustment in the decision process [43]. To understand whether monkeys adjusted their strategy for localized and scattered maps, we estimated the weights of information, reward, and proximity for the scattered maps. We found that the weights of the information map were higher for both monkeys foraging the scattered maps, compared to localized maps (Fig. 5F; compare to Fig. 4D; Fig 5 G), indicating a higher contribution of information-seeking, when foraging an scattered maps.

## Discussion

We investigated the spatial foraging pattern of free-roaming macaques on ambiguous terrain. The foraging terrain consisted of a uniform grid of woodchip piles on the floor, hiding pieces of rewards that were distributed in the grid according to a reward abundance map. Several aspects of our experiment aimed at generating an ambiguous terrain: 1. the environment did not provide any sensory cue to reveal the reward locations 2. Macaques performed the task for a low number of foraging sessions, without prior intense training. This allowed us to understand their foraging strategy in an unfamiliar terrain 3. The type of the abundance map, localized or scattered, was not revealed to the animals using sensory cues. We found that the free-moving monkeys navigate in Lévy-like patterns, search preferably nearby after reward encounters, but do not restrict their search to the vicinity of discovered food. A spatial kernel-based model weighing reward seeking, information seeking, and energy preservation is able to generate diverse ecologically valid foraging pathes. Fitting this model to the experimental data reveals individual differences among four tested animals, as well as adaptation to the different distributions of the hidden food. We will discuss how the observed behaviors mark an ecologically valid solution to seek information when foraging an ambiguous terrain.

Foraging individuals need to continuously evaluate and decide between exploiting the immediately available nearby food options or exploring more remote, often unknown alternatives [21]. We studied the case where a uniform terrain of potential food sources needed to be searched through to find potentially hidden rewards. The macaque’s foraging patterns revealed that they incorporated information-seeking in their strategy when foraging such ambiguous terrain: The distribution of distance traveled between consecutive searches followed a Lévy-like pattern. This means that monkeys occasionally traveled a long distance between consecutive searches, as revealed by a heavy-tailed power probability distribution over the step sizes. In other words, the animals favored the piles in the vicinity of discovered rewards but did not restrict their search by dwelling around them; instead, they continued to explore the entire terrain. We explained information-seeking on our ambiguous terrain using a kernel-based spatial model incorporating a map of information where the unsearched piles, i.e., the information sources, were located.

The observed Lévy-like random walk pattern indicates that the log-likelihood of choosing a pile was inversely proportional to the log distance from this pile. Occasionally choosing to travel a long distance indicates that macaques continuously balanced energy preservation, i.e., crawling the field by choosing nearby piles, with random exploration, i.e., zig-zagging the grid, choosing piles regardless of their distance. This observation is consistent with the search patterns of many species, ranging from microorganisms to human hunter-gatherers, in their natural habitats [27]– [30], [44]–[46]. However, for numerous reasons we do not call the foraging pattern of our macaques a Lévy walk or a Lévy flight: First, the scale-freeness of a Lévy walk was limited by the relatively small size of the terrain, spanning only about two orders of magnitude (30cm to 3m). Second, a property of Lévy walk is the uniform distribution of the heading angle at each step [47]. In our experiment, this assumption was unmet because the heading angle of the animals was discretized due to the finite resolution of the pile grid. Third, even if the terrain was more fine-grided, we speculate that monkeys will likely choose their heading angle non-uniformly, favoring the directions in their field of view. Therefore, even with a Lévy distribution of the step sizes, the search pattern is not necessarily a Lévy walk [47].

Consistent with previous reports on land and marine animals [31], we found that macaques favor searching the vicinity of previous food encounters. This strategy is particularly efficient when the food distribution is localized. In an extreme case, when the food is distributed within one local patch, a forager may switch from roaming the field to dwelling around the location of the discovered food, a greedy strategy [48] to exhaustively exploit the only food source. Switching from roaming to dwelling after the first food encounter has been reported in other species such as *C*.*elegans* [34]–[36] with distinct neural circuits underpinning behavior in each phase. On our localized maps, a forager, who knows a priori or learns quickly that the filled piles were clustered together, optimally roams around until finding a filled pile, then dwells around to visit all filled piles. However, we found no evidence that macaques learned the local structure of the hidden abundance map during their limited exposure to the task (Fig. S2). Instead, we found evidence that they mildly favored the vicinity of the reward location but continued to explore the terrain.

We developed a model to generate and predict spatial foraging patterns using three principles energy preservation, reward-seeking, and information-seeking. Essentially, we attributed a probability of being chosen to each untouched pile based on preset weights for contributions of energy preservation, reward-seeking, and information-seeking. This probability was updated after each pile search, as in a generic reinforcement learning framework in which the agent learns the value of available options by integrating outcomes over time, updates an internal distribution of values, assigned to available options, and the policy which generates choices using the value distribution over the options [12].

The spatial kernel-based model was used for simulating hypothetical agents as well as investigating foraging paths in the experimental data. In the generative model, this probability distribution was used to generate a sequence of choices while updating the probability distribution after each pile search. In the fitted model, the probability distribution parameters were trained to maximize the likelihood of observing the experimental data. Using the generative model, we generated ecologically valid foraging patterns by choosing the weight composition. Using the fitted model, we found differences across individuals and map types. Comparing information weights across four individuals suggests a higher weight for two younger monkeys, Vin and Nat, than for two older monkeys, Luk and Hum (Fig. S4). Age-related decline in exploration is expected in adults [49]. However, for our monkeys, other factors such as social dominance, experience in lab environments, or food preference may as well explain this distinction. Further investigation using a diverse population of monkeys is needed to explain individual differences in balancing information-seeking with reward-seeking. Comparing information weights across two map types, we found a higher information weight for scattered maps, suggesting an adjustment in the foraging strategy, allowing the animals to succeed in finding scattered rewards.

Taken together, we demonstrate how our Exploration Room platform can be used to study ecologically relevant yet experimentally controlled forms of spatial foraging without extensive training of the animals and thereby learn about the weighing of different search-relevant factors in foraging decisions. Our results suggest an explicit role for information seeking, alongside previously considered energy preservation and reward-seeking in macaques’ foraging strategies, an ecologically valid way of foraging ambiguous terrains. We speculate that macaques have evolved this trait to survive in and thrive under the ambiguity of novel terrains.

## Materials and Methods

### Animal use statement

All procedures comply with the European Directive 2010/63/EU and the German Law and have been approved by the regional authorities (Niedersächsisches Landesamt für Verbraucherschutz und Lebensmittelsicherheit (LAVES)) under the permit number 33.19-42502-04-18/2823. Four rhesus monkeys (Macaca mulatta) were used in the study: Vin (10 years old, 7.0 kg), Luk (20 years, 9.0 kg), Hum (16 years, 15.5 kg), and Nat (7 years, 7.5 kg). The animals were group-housed in the animal facility of the German Primate Center in Göttingen in groups of two or three. The animals have an enriched environment consisting of several wooden structures and toys. The home cages of the animal exceed the size regulations by European guidelines and provide access to natural light in an outdoor space. All animals were trained to climb into a primate chair to transfer from the housing facility into the Exploration Room setup.

### Experimental setup

The experimental setup, Exploration Room [38] was a custom-made room with dimensions of 4.6 m (W), 2.5 m (D), and 2.6 m (H). The skeleton was constructed of an aluminum track system (MiniTec, GmbH, Schönenberg-Kübelberg, Germany). The walls and the floor were tiled with white high-pressure laminate (HPL, Kunststoffplattenonline.de). Two doors along the length gave access to humans, and a gated custom-made tunnel on the opposite side gave access to macaques. The ceiling was covered with a metal mesh grid. The room was lit using 8 LED panels just above this mesh grid. The animal’s position and searching of piles were assessed via video recorded from 2-6 Chameleon3 USB3 cameras (FLIR Systems Inc, Wilsonville, Oregon, U.S.) placed strategically to record monkeys’ full body actions from divese angles. Fig. S5 shows the view from two of these cameras, placed at the ceiling in a central position, at an equal distance (∼1.5m) from each other and the short walls of experimental rooms.

### Behavioral training and testing

Each monkey was habituated to the exploration room before recordings. During the habituation sessions, low-density foraging terrains, i.e., terrains with fewer piles spread further from each other and containing a random number of food items, were used to ensure that the animals searched the piles when encountering them. The woodchips were the same kind used in their housing, facilitating the habituation phases. The high-density pile grid with 81 or 108 piles was used for testing only. The localized abundance maps were determined in each session by randomly choosing one of the piles in the grid to be the center of the filled piles, but excluding the centers for which filled piles were truncated by the borders of the grid. The food items were pieces of banana chips, cucumber, radish, or grapes, depending on the preference of the animal. Only one type of food piece was used in each session. When cucumbers were used, we dampened the woodchips with water to release the natural wood smell strong enough to mask the smell of fresh cucumber pieces. The end of the foraging session was determined when the animal waited near the exit to leave the room or roamed around for longer than 5 minutes without searching.

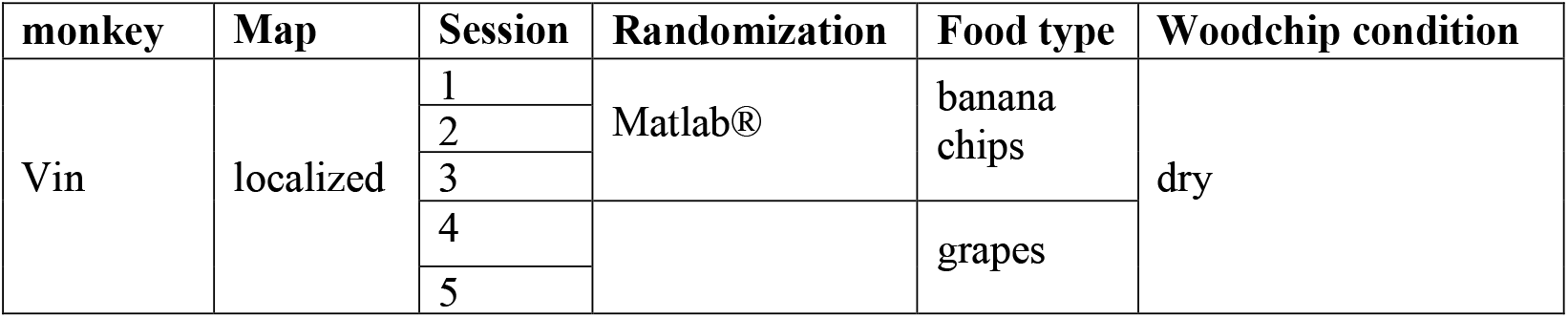
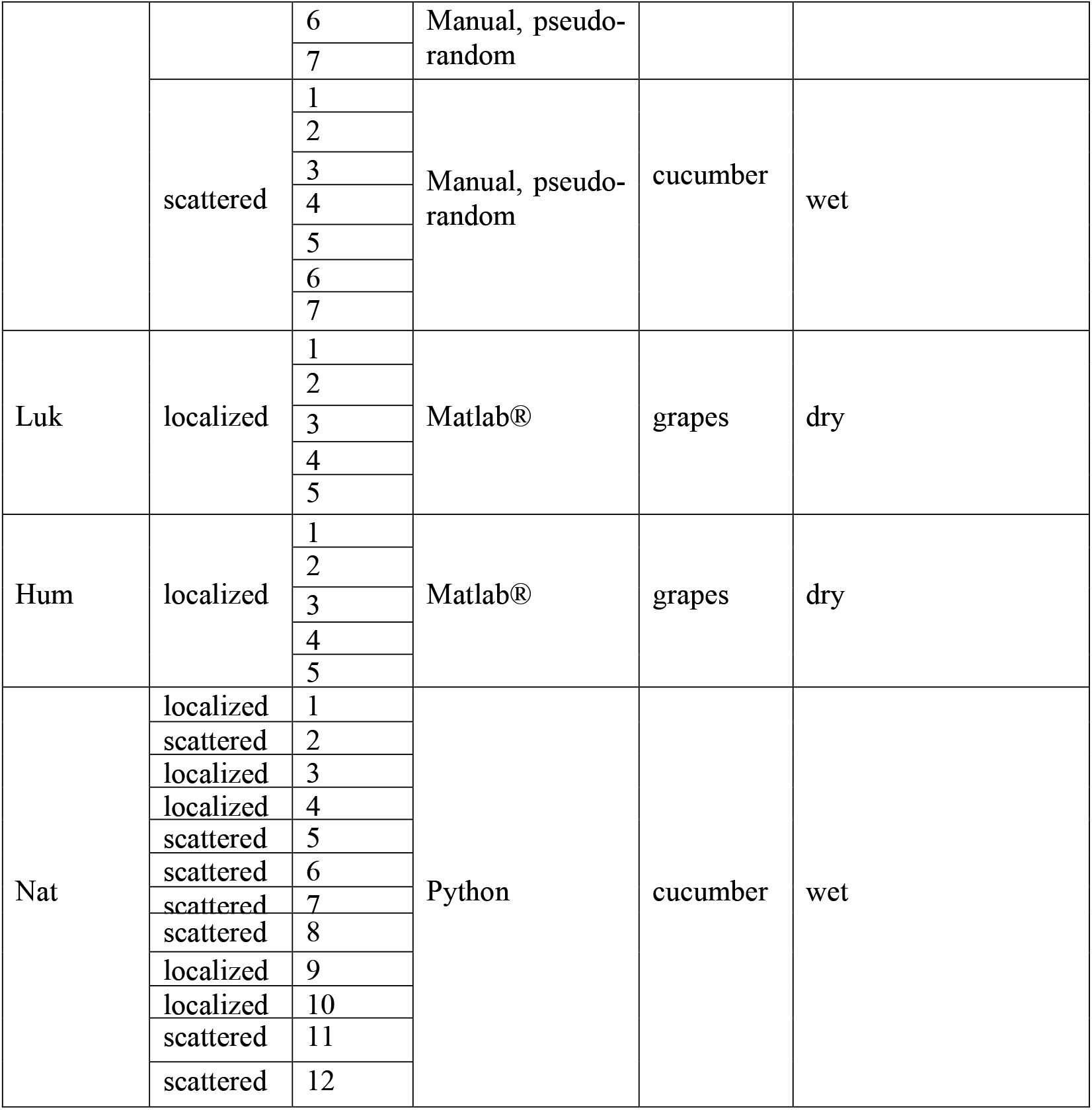

| monkey | Map | Session | Randomization | Food type | Woodchip condition |
| --- | --- | --- | --- | --- | --- |
| Vin | localized | 1 | Matlab® | banana chips | dry |
|  |  | 2 |  |  |  |
|  |  | 3 |  |  |  |
|  |  | 4 |  | grapes |  |
|  |  | 5 |  |  |  |
|  |  | 6 | Manual, pseudo-random |  |  |
|  |  | 7 |  |  |  |
|  | scattered | 1 | Manual, pseudo-random | cucumber | wet |
|  |  | 2 |  |  |  |
|  |  | 3 |  |  |  |
|  |  | 4 |  |  |  |
|  |  | 5 |  |  |  |
|  |  | 6 |  |  |  |
|  |  | 7 |  |  |  |
| Luk | localized | 1 | Matlab® | grapes | dry |
|  |  | 2 |  |  |  |
|  |  | 3 |  |  |  |
|  |  | 4 |  |  |  |
|  |  | 5 |  |  |  |
| Hum | localized | 1 | Matlab® | grapes | dry |
|  |  | 2 |  |  |  |
|  |  | 3 |  |  |  |
|  |  | 4 |  |  |  |
|  |  | 5 |  |  |  |
| Nat | localized | 1 | Python | cucumber | wet |
|  | scattered | 2 |  |  |  |
|  | localized | 3 |  |  |  |
|  | localized | 4 |  |  |  |
|  | scattered | 5 |  |  |  |
|  | scattered | 6 |  |  |  |
|  | scattered | 7 |  |  |  |
|  | scattered | 8 |  |  |  |
|  | localized | 9 |  |  |  |
|  | localized | 10 |  |  |  |
|  | scattered | 11 |  |  |  |
|  | scattered | 12 |  |  |  |

### Action labeling

Pile searches of each monkey were detected using a custom-written, freely available action labeling software [50] or a custom code in Matlab®. A unique label was assigned to each pile, as in Fig. S5. Each pile search was identified using the searched pile’s label and the search’s time. The sequence of the searched piles in each session was subsequently converted into 2D coordinates of the searched location.

### Power function

To determine whether the distribution of the step sizes was Lévy-like, we used the following power function

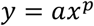

On a double-log scale, this formula transforms to

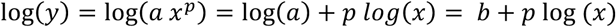

which is a line with a bias *b* and a slope *p*. Accordingly, we fitted linear models to step-sizes in log-log space using a least-square regression using the built-in function ‘regress’ in Matlab®.

### Calculating convolutedness index

The convolutedness of a sub-path was calculated using the angles between the direction of consecutive steps and the length of steps according to

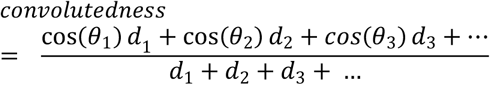

In which d1, d2, … are the length of steps in meters and θ1, θ2, … are the angle between the current step and the previous step, except for θ1, which we use the last step of the path as the step before the first step. The further illustration clarifies the definition of angles between consecutive steps.

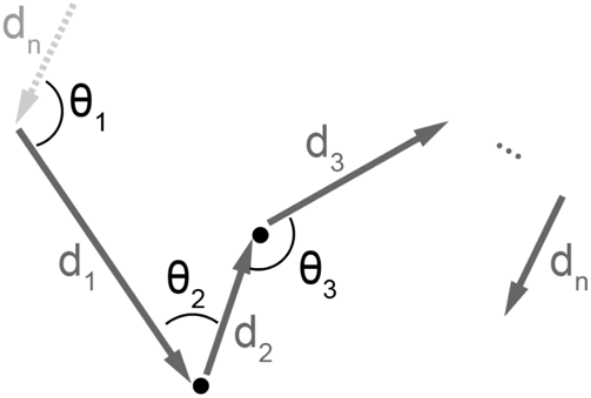

### Generative kernel-based spatial model

The model consisted of a 2-dimensional probability *map*_*prob*_(*x, y*), determining the probability that the simulated agent chooses the pile at the location x and y for the next search. x and y took integer values with each pile representing one unit. The probability map consisted of three maps: the ambiguity map, the reward map, and the proximity map as follow:

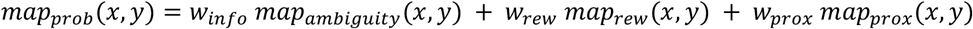

The ambiguity map consisted of a superposition of 2-D Gaussian kernels, each centered at one unsearched pile. Once a pile was searched, the info map was updated by removing the Gaussian kernel associated with that pile. In the generative model of Fig. 4A-C, *σ*_*ambiguity*_, the standard deviation of the ambiguity kernels was 3 piles = 0.9 m.

The reward map started with zeros. After each pile search, if the pile contained rewards, one 2-D Gaussian kernel centered at the searched pile and scaled with the number of rewards was added to the reward map. In the generative model of Fig. 4A-C, *σ*_*rew*_, the standard deviation of the reward kernels was 1 pile = 0.3 m.

The proximity map consisted of one 2-D Gaussian kernel centered at the location of the currently searched pile. In the generative model of Fig. 4A-C, *σ*_*prox*_, the standard deviation of the proximity kernel was 2 piles = 0.6 m.

In the generative model, the next pile was sampled from *map*_*prob*_(*x, y*)^*p*^. When *p* > 10, the generative model was almost deterministic, choosing arg max [*map*_*prob*_(*x, y*)] as the next pile. For simulations in Fig. 4A-C, we used *p* = 5.

### Fitting the kernel-based spatial model to the experimental data

We used a maximum likelihood approach to fit the model to the experimental data. We calculated the *LL*, the log-likelihood of *map*_*prob*_(*w*_*info*_, *w*_*rew*_, *σ*_*info*_, *σ*_*rew*_, *σ*_*prox*_). *w*_*prox*_ was set so that *w*_*info*_ + *w*_*rew*_ + *w*_*prox*_ = 1. *map*_*prob*_ was computed for each foraging step by updating ambiguity, reward, and proximity maps from the previous step and re-normalizing *map*_*prob*_ to have a sum of 1. We chose each of *σ*_*ambiguity*_, *σ*_*rew*_, and *σ*_*prox*_ from a list of 11 values: 0.1, 0.4, 0.7, 1, 1.3, 1.6, 1.9, 3, 4, 5, and 6 (going through all 1331 possible permutations), then used *fmincon* function in Matlab® to find *w*_*info*_, *w*_*rew*_ to minimize −*LL*, constraining the free parameters to [0,1]. We used only one initial set of values because *map*_*prob*_ has only one global maximum in the 2-dimensional space of *w*_*info*_ and *w*_*rew*_. After going through all 1331 permutations of *σ*_*ambiguity*_, *σ*_*rew*_, and *σ*_*prox*_, we chose these 3 parameters to maximize *LL*. For cross-validating the model, we repeated this procedure for 100 random subsets of pile searches for each monkey, each containing 80% of the total number of pile searches.

### Statistical analysis

A two-sided Wilcoxon signed-rank test was used, except where another test was indicated. When multiple data groups were compared, false discovery rate (FDR) correction for multiple comparisons [51] was used to correct the *p*-values.

## Contributions

Conceptualization, N.S., Z.A., Y.B., and A.G.; Methodology, N.S., Z.A., I.L., and A.G.; Investigation, N.S., Z.A., I.L.; Analysis and modeling, N.S., Writing – Original Draft, N.S.; Writing – Review & Editing, N.S., Z.A., Y.B., I.L., A.G.; Funding Acquisition, A.G.; Resources, A.G.; Supervision, N.S. and A.G.

## Acknowledgment

The authors thank Igor Kagan (decision and awareness group, Cognitive Neuroscience lab, Germany Primate Center, Göttingen, Germany) for his constructive feedback on the manuscript. This publication was funded by the Deutsche Forschungsgemeinschaft (DFG, German Research Foundation) -SFB 1528 Cognition of Interaction and the Ministry for Science and Culture of Lower Saxony (MWK, grant ZN3422 DeMoDiag).

## Use of generative AI

Grammarly (Grammarly, Inc.) and ChatGPT 4o (open AI) were used to improve the text’s Grammar, style, and clarity.

## Code and data availability

Custom-written code and accompanying data are available at https://data.goettingen-research-online.de/dataset.xhtml?persistentId=doi:10.25625/DBAN4V

## Notes

### Competing Interest Statement

The authors have declared no competing interest.

### Summary of Updates

An author summary, digestible to non-expert readers, was added a link to a repository containing the codeand the data was added

