## supplemental figures for "Freely foraging macaques value information in ambiguous terrains"

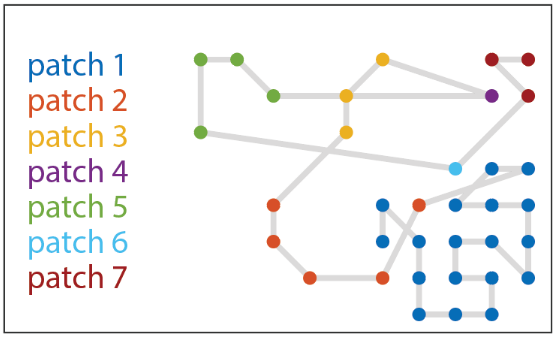


**Figure S1 |** A foraging session of monkey Hum in which the foraging path resembles a patch-wise search. A patch was identified as a set of consecutive searches for which the step size between each search and the previous search is smaller than 1.5 meters. A patch-wise pattern was not consistently observed across other sessions/monkeys. Instead, we found unimodal distributions of step sizes, as in Fig. 2B.

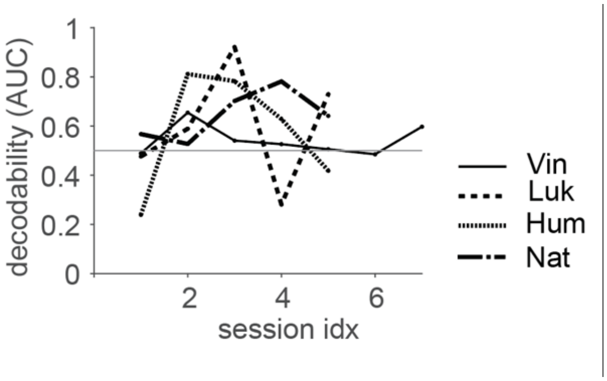


**Figure S2 |** Decodability of two groups of step sizes: after encountering filled and empty piles across sessions of one monkey. Decodability was quantified as the area under the R.O.C. The chance level is 0.5. None of the monkeys showed a linear trend across sessions (p = 0.9 (Vin), 0.8 (Luk), 0.9 (Hum), 0.3 (Nat) for a linear regression model).

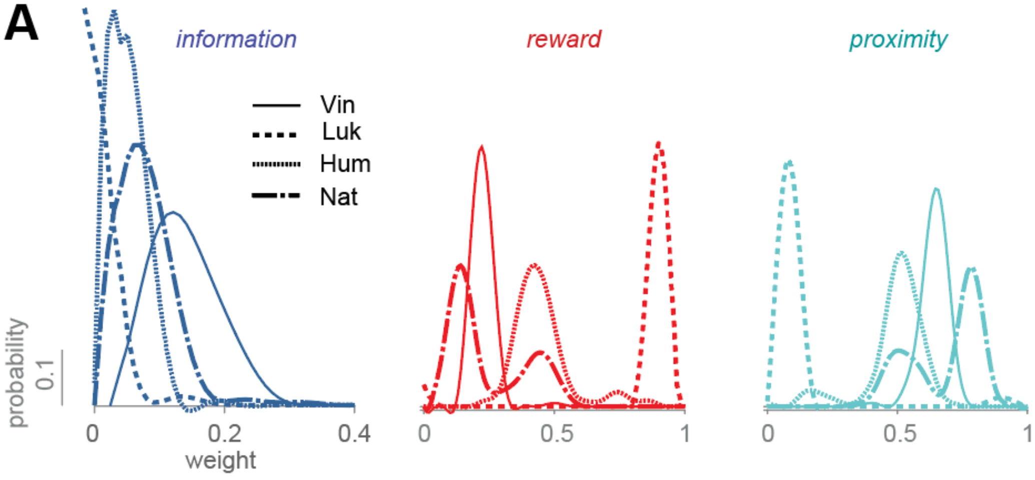


**Figure S3 |** Comparing weights of the spatial model across monkeys or maps. **A)** Comparing the information, reward, and proximity weights across monkeys. Vin weighed information higher than other monkeys (decodability > 0.88) while Luk weighed rewards higher than others (decodability > 0.95).

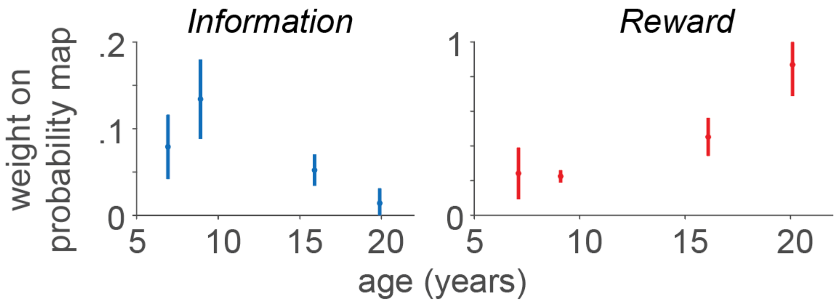


**Figure S4 |** The information and reward weights as a function of monkeys' age.

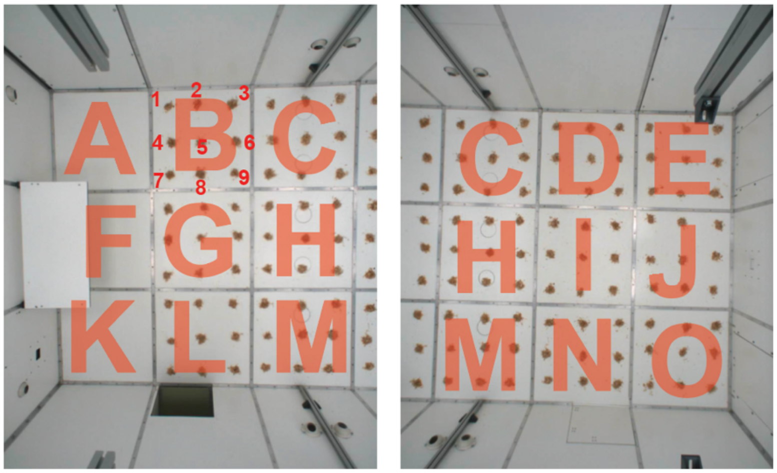


**Figure S5 |** The bird's eye view of the foraging terrain using two overhead cameras, overlapped with a map of identifying labels for each pile. For example, the topmost left pile was B1, and the bottommost right label was O9.
